# Longitudinal plasma metabolomics of aging and sex

**DOI:** 10.1101/436931

**Authors:** Burcu F. Darst, Rebecca L. Koscik, Kirk J. Hogan, Sterling C. Johnson, Corinne D. Engelman

## Abstract

Understanding how metabolites are longitudinally influenced by age and sex could facilitate the identification of metabolomic profiles and trajectories that indicate disease risk. We investigated the metabolomics of age and sex using longitudinal plasma samples from the Wisconsin Registry for Alzheimer’s Prevention (WRAP), a cohort of participants who were dementia free at enrollment. Metabolomic profiles were quantified for 2,316 fasting plasma samples among 1,187 participants, each with up to three study visits. Of 1,097 metabolites tested, 608 (55.4%) were associated with age and 680 (62.0%) with sex after correcting for multiple testing. Approximately twice as many metabolites were associated with age in stratified analyses of women versus men, and 63 metabolite trajectories significantly differed by sex, most notably including sphingolipids, which tended to increase in women and decrease in men with age. Using genome-wide genotyping, we also report the heritabilities of metabolites investigated, which ranged dramatically (0.2-99.2%); however, the median heritability of 36.2% suggests that many metabolites are highly influenced by a complex combination of genomic and environmental influences. These findings offer a more profound description of the aging process and may inform many new hypotheses regarding the role metabolites play in healthy and accelerated aging.

## Introduction

The metabolome represents the functional endpoints of a complex network of biological events, including genomic, epigenomic, transcriptomic, proteomic, and environmental factors (Deidda, Piras, Bassareo, Dessalvi, & Mercuro, 2015). Being the final downstream product, the metabolome is the closest to the phenotype among the biological systems (Horgan & Kenny, 2011), making it particularly relevant to investigate. Age is known to be the single largest risk factor of most prevalent diseases in developed countries (Niccoli & Partridge, 2012). A better understanding of how the metabolome changes with age could further reveal the mechanisms by which age influences disease risk and could facilitate the identification of high-risk metabolomic profiles that are suggestive of the early stages of particular diseases.

Previous studies have provided important evidence that age and sex influence the metabolome (Chaleckis, Murakami, Takada, Kondoh, & Yanagida, 2016; Dunn et al., 2015; Krumsiek et al., 2015; Menni et al., 2013; Mittelstrass et al., 2011; Rist et al., 2017; Yu et al., 2012). While informative, these studies are limited by their cross-sectional designs and the relatively small number of metabolites assessed by most. According to the Human Metabolite Database (HMDB) v4.0, there are an estimated 25,424 blood metabolites (Wishart et al., 2018). However, due to current technical limitations in identifying and quantifying metabolites, most recent studies have only been able to confidently capture ~100-600 of these. A larger panel of metabolites will provide a more comprehensive understanding of the metabolomics of age and sex. Further, in order to assess the metabolomics of aging, it is crucial to use a longitudinal study design that can capture age-related phenomena, particularly due to the high variability of metabolites (Makinen & Ala-Korpela, 2016). Longitudinal assessments also facilitate the examination of metabolite trajectories, which can address important biological questions.

Using longitudinal plasma samples from the Wisconsin Registry for Alzheimer’s Prevention (WRAP), we investigated how a large panel of metabolites is influenced by age and sex, and whether metabolite trajectories vary by sex. To facilitate the interpretation of our results and determine whether identified metabolites are more strongly influenced by genetic or environmental factors, we used genome-wide genotyping data to assess the heritability (*h^2^*) of metabolites.

## Results

### Participants

A total of 1,212 WRAP participants with 2,344 longitudinal fasting plasma samples were available for analyses. At the baseline visit for the current study, participants were 61 years old on average, 69% were female, and 94% were Caucasian (Table 1). Most individuals were unrelated (n=825), but 147 families had >1 individual (family sizes ranged from 1-9 members, with an average of 1.2 individuals per family). Analyses stratified by sex included 838 women and 374 men, who had similar characteristics with the exception of more men taking cholesterol lowering medications than women. Participants each had 1,097 plasma metabolites available for analyses, 347 (31.6%) of which were of unknown chemical structure. Correlations between metabolites were assessed using Pearson r and the first available sample for each individual (*i.e.*, using a cross-sectional approach). Metabolites were largely uncorrelated with each other (Figure S1). Properties of each metabolite, such as biochemical name, super pathway, and sub pathway, are described in Table S1.

**Table 1.** WRAP Participant Characteristics at Baseline for the Current Study. Mean (SD) or N (%).

| Characteristic | Overall<br>(N=1,212,<br>obs=2,344) | Male<br>(n=374,<br>obs=731) | Female<br>(n=838,<br>obs=1,613) |
| --- | --- | --- | --- |
| Age (years) | 60.8 (6.7) | 61.2 (6.9) | 60.7 (6.6) |
| Caucasian | 1,135 (93.7) | 351 (93.9) | 784 (93.6) |
| Cholesterol lowering<br>medication | 387 (31.9) | 146 (39.0)* | 241 (28.8)* |
| Sample storage (days) | 1,510.5 (415.7) | 1,511.2 (424.3) | 1,510.2 (412.0) |

obs=number of longitudinal observations
\*Differs between men and women with $P=3.9\text{e-}4$

### Metabolome-wide association study

#### Aging Metabolomics

Associations were tested using linear mixed effects regression models implemented in the SAS MIXED procedure. Primary predictors included age and sex, which were assessed within the same models. To examine effect modification of the metabolomics trajectories by sex, analyses were repeated stratifying the sample by sex. To assess the statistical significance of the effect modification, separate models were run that included an interaction term for age-by-sex using the full sample (men and women combined). All models included random intercepts for within-subject correlations (due to repeated measures) and within-family correlations (due to siblings). Models included fixed effects for age, sex, self-reported race, and cholesterol lowering medication use, which was the most commonly used class of medications in our sample. Sensitivity analyses were performed with an additional fixed effect for plasma sample storage time. Each set of analyses was corrected for multiple testing using the Benjamini-Hochberg (Benjamini & Hochberg, 1995) adjustment with an alpha of 0.05.

All metabolome-wide association results are summarized in Table 2 and detailed in Table S1. After adjusting for multiple testing, the levels of 637 metabolites (58.1% of metabolites assessed) significantly changed with age, 516 of which increased with age (Figure S2A and Figure 1). Of the total 34 steroid lipids tested, 28 significantly decreased with age (including 20/22 androgenic, 5/5 progestin, 4/4 pregnenolone, and 1/3 corticosteroids), while two, 11- ketoetiocholanolone glucuronide, an androgenic steroid, and cortisol, significantly increased with age.

**Table 2.**
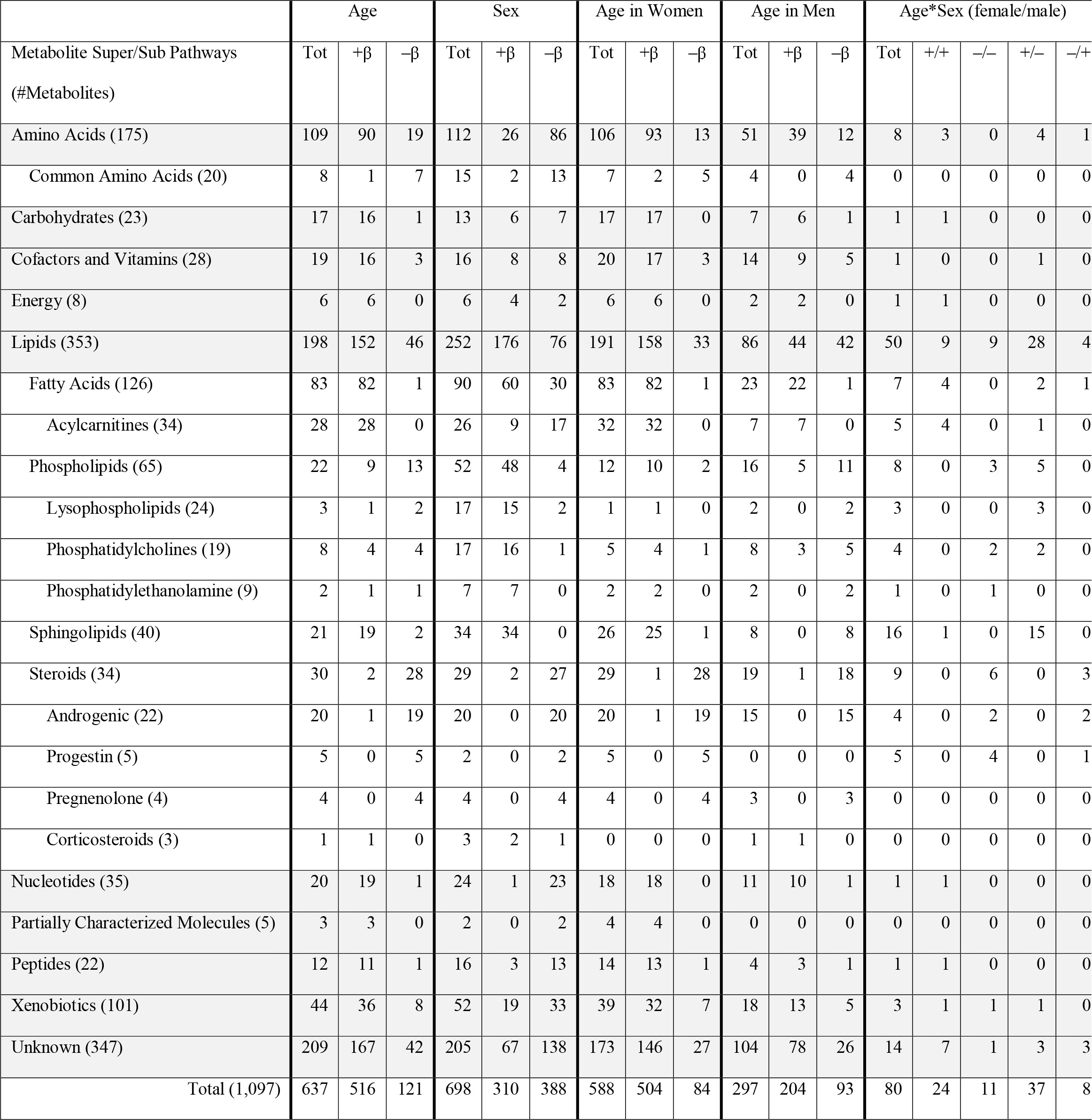
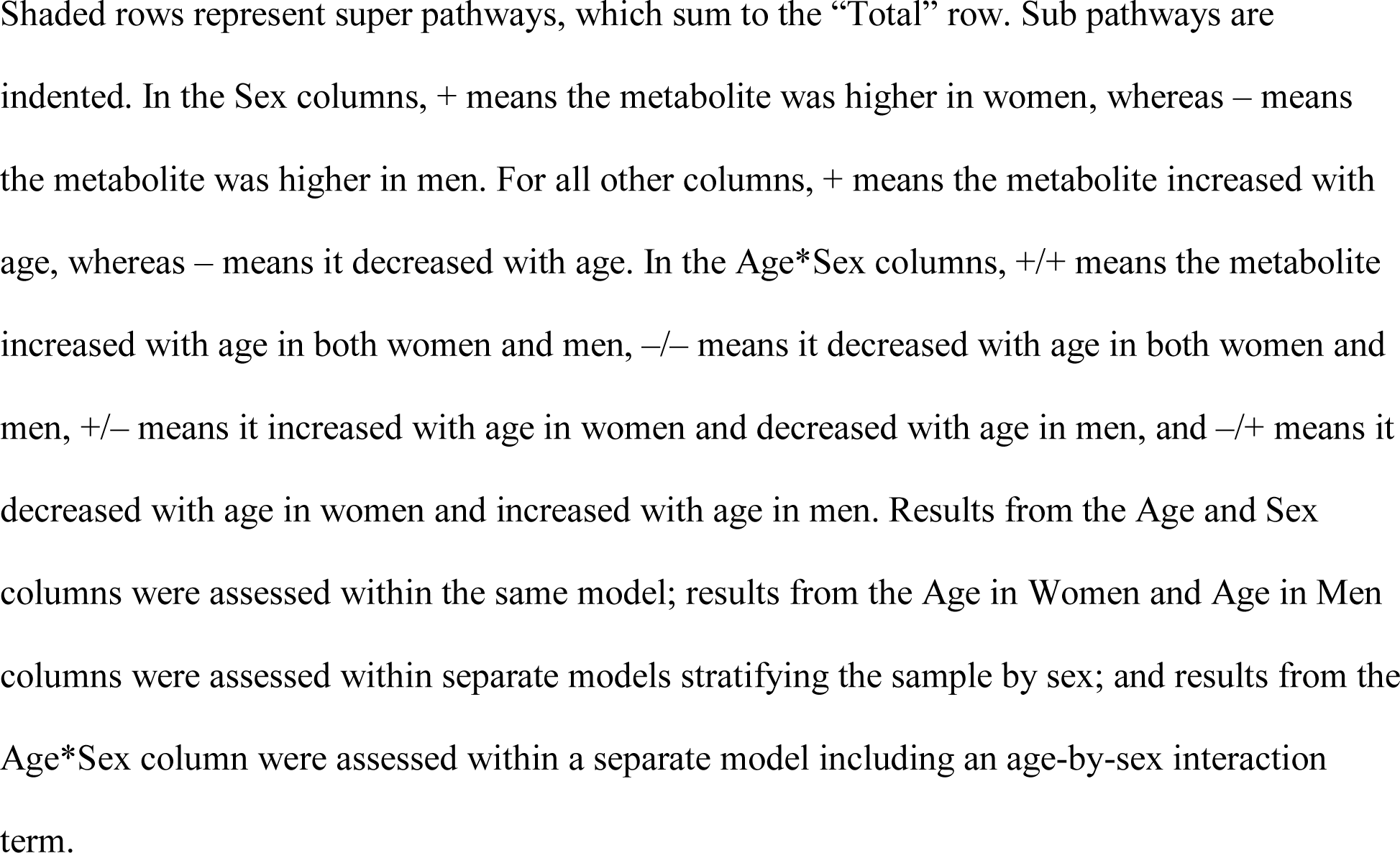
Metabolome-wide association results summary. Number of metabolites associated with each trait by pathways and recurrent sub pathways after correcting for multiple comparisons.

| Metabolite Super/Sub Pathways<br>(#Metabolites) | Age |  |  | Sex |  |  | Age in Women |  |  | Age in Men |  |  | Age*Sex (female/male) |  |  |  |  |
| --- | --- | --- | --- | --- | --- | --- | --- | --- | --- | --- | --- | --- | --- | --- | --- | --- | --- |
| | Tot | +$\beta$ | -β | Tot | +$\beta$ | -β | Tot | +$\beta$ | -β | Tot | +$\beta$ | -β | Tot | +/+ | -/- | +/- | -/+ |
| Amino Acids (175) | 109 | 90 | 19 | 112 | 26 | 86 | 106 | 93 | 13 | 51 | 39 | 12 | 8 | 3 | 0 | 4 | 1 |
| Common Amino Acids (20) | 8 | 1 | 7 | 15 | 2 | 13 | 7 | 2 | 5 | 4 | 0 | 4 | 0 | 0 | 0 | 0 | 0 |
| Carbohydrates (23) | 17 | 16 | 1 | 13 | 6 | 7 | 17 | 17 | 0 | 7 | 6 | 1 | 1 | 1 | 0 | 0 | 0 |
| Cofactors and Vitamins (28) | 19 | 16 | 3 | 16 | 8 | 8 | 20 | 17 | 3 | 14 | 9 | 5 | 1 | 0 | 0 | 1 | 0 |
| Energy (8) | 6 | 6 | 0 | 6 | 4 | 2 | 6 | 6 | 0 | 2 | 2 | 0 | 1 | 1 | 0 | 0 | 0 |
| Lipids (353) | 198 | 152 | 46 | 252 | 176 | 76 | 191 | 158 | 33 | 86 | 44 | 42 | 50 | 9 | 9 | 28 | 4 |
| Fatty Acids (126) | 83 | 82 | 1 | 90 | 60 | 30 | 83 | 82 | 1 | 23 | 22 | 1 | 7 | 4 | 0 | 2 | 1 |
| Acylcarnitines (34) | 28 | 28 | 0 | 26 | 9 | 17 | 32 | 32 | 0 | 7 | 7 | 0 | 5 | 4 | 0 | 1 | 0 |
| Phospholipids (65) | 22 | 9 | 13 | 52 | 48 | 4 | 12 | 10 | 2 | 16 | 5 | 11 | 8 | 0 | 3 | 5 | 0 |
| Lysophospholipids (24) | 3 | 1 | 2 | 17 | 15 | 2 | 1 | 1 | 0 | 2 | 0 | 2 | 3 | 0 | 0 | 3 | 0 |
| Phosphatidylcholines (19) | 8 | 4 | 4 | 17 | 16 | 1 | 5 | 4 | 1 | 8 | 3 | 5 | 4 | 0 | 2 | 2 | 0 |
| Phosphatidylethanolamine (9) | 2 | 1 | 1 | 7 | 7 | 0 | 2 | 2 | 0 | 2 | 0 | 2 | 1 | 0 | 1 | 0 | 0 |
| Sphingolipids (40) | 21 | 19 | 2 | 34 | 34 | 0 | 26 | 25 | 1 | 8 | 0 | 8 | 16 | 1 | 0 | 15 | 0 |
| Steroids (34) | 30 | 2 | 28 | 29 | 2 | 27 | 29 | 1 | 28 | 19 | 1 | 18 | 9 | 0 | 6 | 0 | 3 |
| Androgenic (22) | 20 | 1 | 19 | 20 | 0 | 20 | 20 | 1 | 19 | 15 | 0 | 15 | 4 | 0 | 2 | 0 | 2 |
| Progestin (5) | 5 | 0 | 5 | 2 | 0 | 2 | 5 | 0 | 5 | 0 | 0 | 0 | 5 | 0 | 4 | 0 | 1 |
| Pregnenolone (4) | 4 | 0 | 4 | 4 | 0 | 4 | 4 | 0 | 4 | 3 | 0 | 3 | 0 | 0 | 0 | 0 | 0 |
| Corticosteroids (3) | 1 | 1 | 0 | 3 | 2 | 1 | 0 | 0 | 0 | 1 | 1 | 0 | 0 | 0 | 0 | 0 | 0 |
| Nucleotides (35) | 20 | 19 | 1 | 24 | 1 | 23 | 18 | 18 | 0 | 11 | 10 | 1 | 1 | 1 | 0 | 0 | 0 |
| Partially Characterized Molecules (5) | 3 | 3 | 0 | 2 | 0 | 2 | 4 | 4 | 0 | 0 | 0 | 0 | 0 | 0 | 0 | 0 | 0 |
| Peptides (22) | 12 | 11 | 1 | 16 | 3 | 13 | 14 | 13 | 1 | 4 | 3 | 1 | 1 | 1 | 0 | 0 | 0 |
| Xenobiotics (101) | 44 | 36 | 8 | 52 | 19 | 33 | 39 | 32 | 7 | 18 | 13 | 5 | 3 | 1 | 1 | 1 | 0 |
| Unknown (347) | 209 | 167 | 42 | 205 | 67 | 138 | 173 | 146 | 27 | 104 | 78 | 26 | 14 | 7 | 1 | 3 | 3 |
| Total (1,097) | 637 | 516 | 121 | 698 | 310 | 388 | 588 | 504 | 84 | 297 | 204 | 93 | 80 | 24 | 11 | 37 | 8 |

Shaded rows represent super pathways, which sum to the “Total” row. Sub pathways are indented. In the Sex columns, + means the metabolite was higher in women, whereas – means the metabolite was higher in men. For all other columns, + means the metabolite increased with age, whereas – means it decreased with age. In the Age\*Sex columns, ++ means the metabolite increased with age in both women and men, -- means it decreased with age in both women and men, +/- means it increased with age in women and decreased with age in men, and -/+ means it decreased with age in women and increased with age in men. Results from the Age and Sex columns were assessed within the same model; results from the Age in Women and Age in Men columns were assessed within separate models stratifying the sample by sex; and results from the Age\*Sex column were assessed within a separate model including an age-by-sex interaction term.

**Figure 1.**
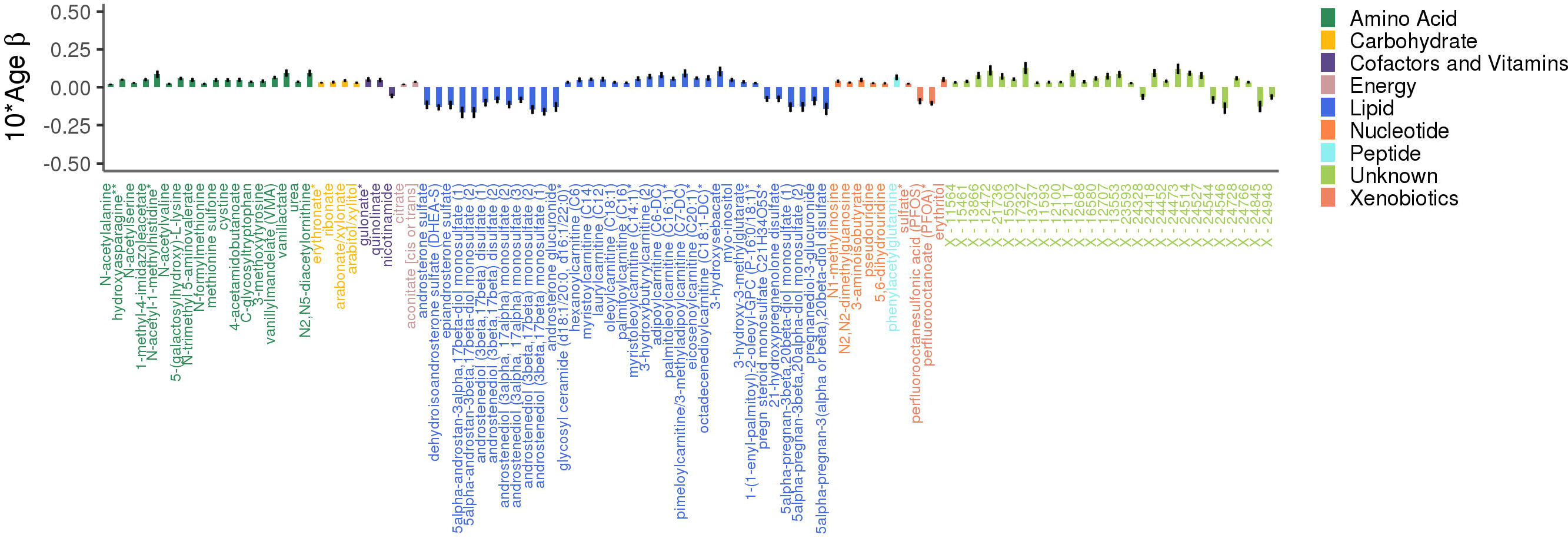
Adjusted effects of a 10-year increase in age on the top 100 metabolites most strongly influenced by age. Positive values indicate the amount a metabolite increased over 10 years, whereas negative values indicate the amount a metabolite decreased over 10 years. Black vertical lines indicate 10*standard errors.

Higher levels of most fatty acid lipids were associated with increased age (including 13/14 long chain fatty acids, 28/34 acylcarnitines, and 41/78 other fatty acids), with the exception of eicosanodioate (C20-DC), a dicarboxylate fatty acid that decreased with age. Higher levels of sphingolipids tended to be associated with increased age (19/21 associated sphingolipids).

The majority of amino acids associated with age increased with age (82.6% or 90/109 associated amino acids), including glutamine, one of the 20 common amino acids that are encoded directly by the genetic code. Seven other common amino acids decreased with age (histidine, threonine, tryptophan, leucine, methionine, aspartate, and asparagine), while the 12 others were not associated with age.

#### Sex Metabolomics

Six hundred and ninety-eight metabolites (63.6% of metabolites assessed) significantly differed by sex, with the slight majority (388 metabolites or 55.6%) found in lower levels in women (Figure 2B and Figure 2). Of the metabolites associated with sex, 415 were also associated with age. Twenty-nine steroid lipids were associated with sex, all of which were found in significantly lower levels in women, with the exception of two corticosteroids (cortisol and corticosterone), which were found in higher levels in women. Androgenic steroids constituted the three metabolites most strongly associated with sex (5alpha-androstan-3alpha, 17beta-diol monosulfate, *P*<4.0e-308, 5alpha-androstan-3alpha, 17beta-diol 17-glucuronide, *P*=4.3e-226, and 5alpha-androstan-3alpha, 17beta-diol disulfate, *P*=2.6e-185).

**Figure 2.**
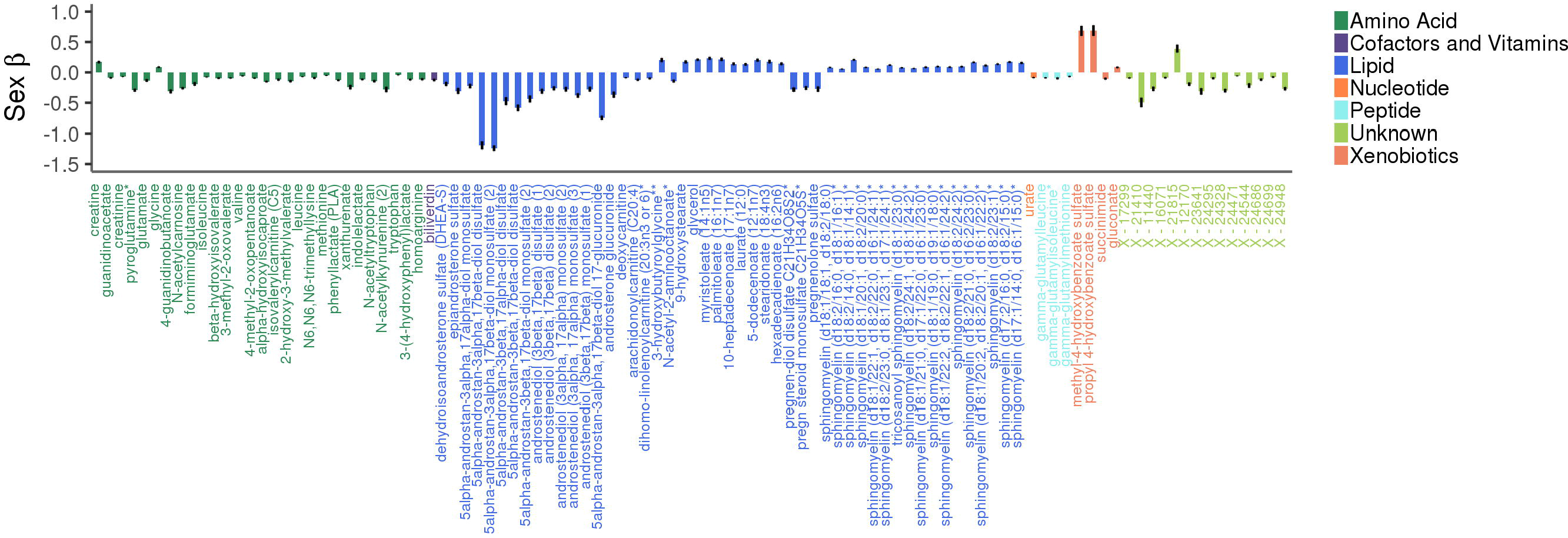
Adjusted effects of the top 100 metabolites most strongly influenced by sex. Positive values indicate that the metabolite was higher in women, whereas negative values indicate that the metabolite was higher in men. Black vertical lines indicate 10*standard errors.

Ninety fatty acids were associated with sex, 60 of which were found in higher levels in women. Acylcarnitine fatty acids were an exception, as 17/26 significantly associated acylcarnitines were found in lower levels in women. Among all tested phospholipids, 73.8% (48/65) were higher in women, as were 85% (34/40) of all tested sphingolipids.

The majority of amino acids associated with sex were found in lower levels in women (76.8% or 86/112), including 13 of the 20 common amino acids (alanine, tyrosine, methionine, arginine, proline, aspartate, asparagine, tryptophan, glutamate, phenylalanine, and the three branched-chain amino acids (BCAAs): leucine, isoleucine, and valine), while two were found in higher levels in women (glycine and serine). The remaining five did not significantly differ by sex.

#### Effect Modification of Metabolomics Trajectories by Sex

Analyses stratified by sex identified 588 metabolites (53.6% of metabolites assessed) that were significantly associated with age among women (Figure S3A and Figure S4) and 297 metabolites (27.1% of metabolites assessed) among men (Figure S3B and Figure S5), with 206 being common to both groups.

The trajectories of 80 metabolites (7.3%) significantly differed over time by sex (Figures S3C and S6). Of the four most significant metabolites, three were sphingolipids, which were also the largest group of metabolites whose trajectories differed by sex (20.0% or 16/80). Fifteen of these sphingolipids increased with age among women and decreased with age among men.

Several other groups of metabolites had trajectories that also differed by sex, including seven fatty acids, six of which showed larger increases with age among women than men; nine steroid lipids, eight of which showed larger decreases with age among women than men; eight phospholipids, five of which increased in women and decreased in men with age; and cholesterol, which increased in women and decreased in men with age.

### Metabolite heritability estimates

The *h^2^* of each metabolite was estimated using a variance components method that jointly models narrow-sense *h^2^* and the *h^2^* explained by genotyped variants (Zaitlen et al., 2013), which allows for the inclusion of both closely and distantly related individuals, as implemented in GCTA (Yang, Lee, Goddard, & Visscher, 2011). A genetic relationship matrix was created from 272,839 weakly linked (R^2^<0.50) and common (MAF>0.05) directly genotyped variants. Analyses of *h^2^* were cross-sectional, using the first available metabolomics sample for 1,111 Caucasians that had both metabolomic and genomic data, and adjusted for sex and age. To assess whether metabolite *h^2^* could influence the effect of age or sex on metabolite levels, Pearson r was used to calculate correlations between *h^2^* estimates and the strength of associations (*i.e.*, *P* - values) for age and sex.

Metabolite *h^2^* estimates ranged widely (0.2-99.1%) and had a median *h^2^* of 36.3%, with a first quartile (Q_1_) of 25.5% and a third quartile (Q_3_) of 49.7% (Figure 3 and Table S1). The metabolites with the lowest *h^2^* were three lipids: adipoylcarnitine (C6-DC), an acylcarnitine (*h^2^* = 0.2%), 15-methylpalmitate (i17:0), a branched fatty acid (*h^2^*=0.2%), and glycosyl-N-stearoyl-sphingosine (d18:1/18:0), a ceramide lipid (*h^2^*=0.6%). The metabolites with the highest *h^2^* estimates were two unknown metabolites (X-12093 and X-24328, *h^2^*=99.1% and 91.1%, respectively) and a nucleotide involved in purine metabolism (N2,N2-dimethylguanosine, *h^2^*=90.0%). Metabolon recently identified X-12093 as N2-acetyl, N6 methyllysine, an amino acid in the lysine catabolic sub pathway.

**Figure 3.**
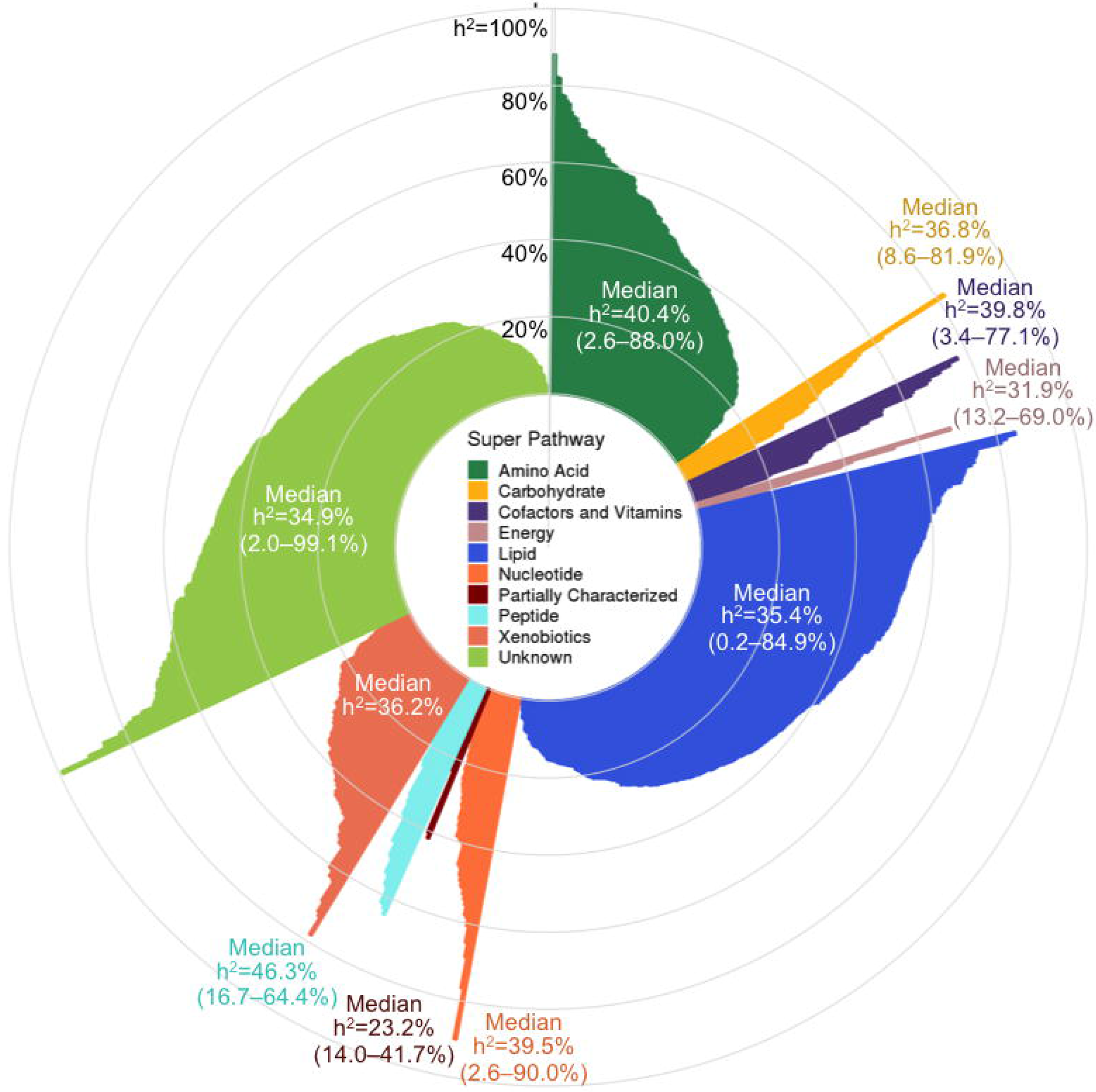
Pinwheel plot of metabolite heritabilities. Each bar indicates the heritability of the corresponding metabolite. Metabolite names are indicated in the outer circle.

Super pathway median *h^2^* estimates ranged from 23.2-46.3%, with peptides having the highest median, followed by amino acids (40.4%), and partially characterized molecules having the lowest median, although the latter pathway only contained five metabolites. Among the metabolite subgroups that were recurrent themes in our association results (*i.e.*, sub pathways highlighted in Table 2), the 20 common amino acids had a median *h^2^* of 49.3% (Q_1_-Q_3_: 36.9-65.1%); fatty acids overall had a median *h^2^* of 30.3% (Q_1_-Q_3_: 16.9-42.4%), while acylcarnitines had a slightly higher median *h^2^* of 41.3% (Q_1_-Q_3_: 26.6-56.0%); phospholipids overall had a median *h^2^* of 35.9 (Q_1_-Q_3_: 24.7-53.3%), while phosphatidylcholines had a slightly lower median *h^2^* of 30.6% (Q_1_-Q_3_: 22.6-39.6%); sphingolipids had a median *h^2^* of 42.0% (Q_1_-Q_3_: 31.8- 51.7%); and steroid lipids overall had a median *h^2^* of 39.6% (Q_1_-Q_3_: 35.0-50.7%), while androgenic steroids had a median *h^2^* of 42.5% (Q_1_-Q_3_: 37.6-50.7%).

Metabolites associated with age and sex had *h^2^* estimates that were representative of overall metabolite *h^2^* estimates. Among the 608 metabolites associated with age, the median *h^2^*=36.1% and Q_1_-Q_3_: 25.9-50.0%. Similarly, among the 680 metabolites associated with sex, the median *h^2^*=37.1% and Q_1_-Q_3_: 26.0-50.6%. Overall, metabolite *h^2^* estimates were not correlated with the strength of associations for age or sex (Pearson r=-0.03 and -0.02, respectively).

## Discussion

To our knowledge, this is the first longitudinal metabolomics assessment of aging and sex and uses one of the largest panels of metabolites reported to date. Our results provide strong evidence that most plasma metabolite levels are highly influenced by aging and that aging has a broader effect on metabolites in women than men. Metabolites are also highly influenced by sex, with men and women having substantially different metabolomic profiles. We report *h^2^* estimates on more metabolites than previously reported and find that the variation of only a few metabolite levels can be attributed almost entirely to either genetic or environmental influences. Rather, most are influenced by a complex combination of genetic and environmental factors, consistent with previous studies (Long et al., 2017; Shin et al., 2014). How heritable a metabolite was did not appear to influence the effect of age or sex on metabolite levels.

Differences in levels of plasma lipid steroids, including androgens, progestins, and pregnenolones, were among the most significant findings for both age and sex. The steroid sex differences serve as a proof of concept, as it is well established that androgens are present in lower levels in women than men (Goodman-Gruen & Barrett-Connor, 2000). Androgens are also known to decrease with age among men in both plasma (Ferrini & Barrett-Connor, 1998) and serum (Harman et al., 2001), and also decline with age in serum among women (Spencer, Klein, Kumar, & Azziz, 2007), perhaps most steeply during early reproductive years (Davison, Bell, Donath, Montalto, & Davis, 2005).

The plasma metabolites we identified to be associated with sex and age are consistent with findings from previous cross-sectional studies. The UK Adult Twin Registry (TwinsUK) study reported 165 out of 280 (58.9%) tested serum and plasma metabolites to be associated with age in cross-sectional analyses (Menni et al., 2013). Our data had 114 of these 165 metabolites, of which 71 were significantly associated with age, and 65 had effects that were in the same direction as those reported in the TwinsUK study (Table S2). The metabolites that had the opposite direction of effect between studies were four amino acids (dimethylarginine, leucine, asparagine, and tryptophan), one nucleotide (uridine), and one xenobiotic (theophylline), all of which we reported decreased with age, with the exception of dimethylarginine, which increased with age, contradictory to findings from the TwinsUK study. However, other studies have reported that serum tryptophan levels decrease with age (Dunn et al., 2015; Yu et al., 2012). Among the 65 metabolites with the same effect, 29 were lipids, all of which increased with age (the majority were fatty acids, including 10 long chain fatty acids, six polyunsaturated fatty acids, and six other fatty acids), and 13 were amino acids (including glutamine, which increased with age, and histidine and aspartate, which both decreased with age).

The Cooperative Health Research in the Region of Augsburg (KORA F4) study, which was also cross-sectional, reported 180 out of 507 (35.5%) tested serum metabolites to be associated with sex (Krumsiek et al., 2015). Our data had 98 of these 180 metabolites, of which 84 were significantly associated with sex, and all had effects that were in the same direction as those reported in KORA F4 (Table S3). Among these were 33 amino acids (including 11 common amino acids, all of which were lower in women except glycine and serine, which is also consistent with Mittelstrass et al. (Mittelstrass et al., 2011)); 18 lipids (including five long chain fatty acids and three medium chain acids, all of which were higher in women, and three androgenic steroids, all of which were lower in women); and 18 unknown metabolites (all but one were lower in women). The single most significant finding in the KORA F4 study was the third most significant in our study (5alpha-androsta-3beta, 17beta-diol disulfate, an androgenic steroid; the two other androgenic steroids that were our first and second most significant sex findings were not tested in the KORA F4 study). Also consistent with our findings, other studies have reported serum and plasma phosphatidylcholines and sphingolipids levels to be higher in women than men (Gonzalez-Covarrubias et al., 2013; Mittelstrass et al., 2011; Rist et al., 2017), and serum acylcarnitines to be lower in women (Mittelstrass et al., 2011).

Consistent with results from our sex-stratified analyses, a previous KORA F4 publication also reported serum sphingolipids to increase in concentrations with age among women and acylcarnitines to increase with age among both women and men (Yu et al., 2012). The KORA F4 study, which had a sample of 1,038 women and 1,124 men, also similarly found twice as many metabolites associated with age among women than men. This suggests that our similar observation may not be driven solely by the differences in sample sizes between women and men in our study and that it may have biological implications; *i.e.*, aging may influence a wider breadth of metabolites in women than men. A probable cause for such a difference may be that during menopause, women experience very abrupt and dramatic hormone changes and loss of ovarian function, whereas during “andropause”, men experience a gradual loss of hormones and decline in fertility(Vermeulen, 2000). These hormonal changes could be associated with other metabolic changes as well. Post-menopausal women have higher levels of sphingomyelins, fatty acids, acylcarnitines, lysophosphatidylcholines, and several amino acids than pre-menopausal women (Auro et al., 2014; Ke et al., 2015), and a recent study found that plasma and urine metabolomics can be used to predict menopause status with 90% accuracy. Moreover, androgenic steroids have been linked to lipid levels in postmenopausal women (Noyan, Yucel, & Sagsoz, 2004). Given that the baseline average ages of women and men in our sample are each ~61 years old, it is likely that our results are indicative of hormonal changes that occur in later ages and that most of our female participants have undergone menopause. It will be crucial to replicate these findings with a metabolomics panel that captures a larger proportion of the ~25,000 known blood metabolites in order to determine the validity of this hypothesis.

Among the 80 metabolites with different trajectories between women and men were sphingolipids, phosphatidylcholines, and cholesterol. Metabolites from the latter two subgroups have been previously reported to have similar trajectories as what we identified, *i.e.*, increasing with age in women and decreasing in men (Auro et al., 2014). To our knowledge, a decrease of sphingolipid levels in men as our results suggest has not been previously reported. However, greater sphingomyelin increases with age in women than men have been previously described(Mielke et al., 2015).

We compared our metabolite *h^2^* estimates to those recently estimated from a twin study of 1,930 individuals in the TwinsUK cohort (Long et al., 2017). Among the 466 metabolites overlapping with our study, *h^2^* estimates were only moderately correlated (Pearson r=0.36) and our estimates were 9.6 percentage points lower on average. However, our metabolite *h^2^* estimates were 8.9 percentage points higher on average (and had a lower correlation of r=0.25) when comparing 191 overlapping metabolite *h^2^* estimates from an earlier twin study based on 7,824 individuals from both the KORA F4 and TwinsUK cohorts (Shin et al., 2014). Interestingly, despite having some overlapping participants, *h^2^* estimates between these two previous studies were only moderately correlated: among 163 overlapping metabolite *h^2^* estimates, Pearson r was 0.38, with estimates based on the TwinsUK cohort being 18.8 percentage points higher on average than the combined KORA and TwinsUK study. Differences in *h^2^* estimates may be driven by differences in population composition and size, phenotypic variation, and analytic approaches.

This study was not without limitations. Our findings are likely driven by our panel of metabolites, and it is possible that a different panel of metabolites could produce different results. Many of our findings are in accordance with previous publications, thereby strengthening confidence in our results that have not been previously investigated with regards to age and sex. Accordingly, it will be crucial to replicate novel findings with an external cohort. However, we also identified several inconsistencies between our study and others regarding *h^2^* estimates and a few of our association results, which could have been due to differences in study designs and sample populations. This challenge is common (Enche Ady et al., 2017), as the field of metabolomics is rapidly developing and widely accepted standards for quality control techniques are forthcoming. Differences in platforms, quantification techniques, statistical analysis methods, laboratory techniques for sample handling (*i.e.*, anti-coagulation method, preservation, storage duration), and fasting status at the time of the sample draw may result in large variations from one study to another (Gonzalez-Dominguez, Sayago, & Fernandez-Recamales, 2017). The metabolomics quality control process we have outlined here as well as that described in Voyle et al. (Voyle et al., 2016) could serve as guidelines for future studies. Many of our findings included metabolites that had unknown chemical structures, which is a current limitation of the field of metabolomics, as it can be difficult and costly to accurately identify metabolites. Further, we only investigated linear effects of age, but non-linear age effects may exist and should be investigated in future investigations.

Using a large panel of longitudinal metabolomics data, we conducted a comprehensive investigation of the influence of aging and sex on metabolomics. Our findings suggest that levels of most metabolites are highly influenced by sex and age, and that sex differentially influences levels and trajectories of many metabolites. These findings underscore the importance of incorporating age and sex in the design and analysis of metabolomics investigations. We also report that many metabolite levels are influenced by a complex combination of both genomic and environmental influences. These findings offer a deeper understanding of the aging process and could inform many novel hypotheses regarding the role of metabolites in healthy and accelerated aging.

## Methods

### Participants

Study participants were from WRAP, a longitudinal study of initially dementia free middle-aged adults that allows for the enrollment of siblings and is enriched for a parental history of Alzheimer’s disease. Further details of the study design and methods used have been previously described (Johnson et al., 2018; Sager, Hermann, & La Rue, 2005). For the current analyses, follow-up occurred every two years. This study was conducted with the approval of the University of Wisconsin Institutional Review Board and all subjects provided signed informed consent before participation.

### Plasma collection and sample handling

Fasting blood samples for this study were drawn the morning of each study visit. Blood was collected in 10 mL ethylenediaminetetraacetic acid (EDTA) vacutainer tubes. They were immediately placed on ice, and then centrifuged at 3000 revolutions per minute for 15 minutes at room temperature. Plasma was pipetted off within one hour of collection. Plasma samples were aliquoted into 1.0 mL polypropylene cryovials and placed in -80°C freezers within 30 minutes of separation. Samples were never thawed before being shipped overnight on dry ice to Metabolon, Inc. (Durham, NC), where they were again stored in -80°C freezers and thawed once before testing.

### Metabolomic profiling and quality control

An untargeted plasma metabolomics analysis was performed by Metabolon, Inc. using Ultrahigh Performance Liquid Chromatography-Tandom Mass Spectrometry (UPLC-MS/MS). Quantification was performed as previously described (Evans et al., 2014); details are outlined in the Supplemental Note. Metabolites within nine super pathways were identified: amino acids, carbohydrates, cofactors and vitamins, energy, lipids, nucleotides, partially characterized molecules, peptides, and xenobiotics.

Up to three longitudinal plasma samples were available for each participant. Metabolites with an interquartile range of zero (*i.e.*, those with very low or no variability) were excluded from analyses (n=178 metabolites). After removing these metabolites, samples were missing a median of 11.7% metabolites, while metabolites were missing in a median of 1.2% of samples. Missing metabolite values were imputed to the lowest level of detection for each metabolite. Metabolite values were median-scaled and log-transformed to normalize metabolite distributions (van den Berg, Hoefsloot, Westerhuis, Smilde, & van der Werf, 2006). If a participant reported that they did not fast or withhold medications and caffeine for at least eight hours, the sample was excluded from analyses (n=159 samples). A total of 1,097 metabolites among 2,344 samples remained for analyses.

### DNA collection and genomics quality control

DNA was extracted from whole blood samples using the PUREGENE^®^ DNA Isolation Kit (Gentra Systems, Inc., Minneapolis, MN). DNA concentrations were quantified using the Invitrogen™ Quant-iT™ PicoGreen™ dsDNA Assay Kit (Thermo Fisher Scientific, Inc., Hampton, NH) analyzed on the Synergy 2 Multi-Detection Microplate Reader (Biotek Instruments, Inc., Winooski, VT). Samples were diluted to 50 ng/ul following quantification.

A total of 1,340 samples were genotyped using the Illumina Multi-Ethnic Genotyping Array at the University of Wisconsin Biotechnology Center (Figure S7). Thirty-six blinded duplicate samples were used to calculate a concordance rate of 99.99%, and discordant genotypes were set to missing. Sixteen samples missing >5% of variants were excluded, while 35,105 variants missing in >5% of individuals were excluded. No samples were removed due to outlying heterozygosity. Six samples were excluded due to inconsistencies between self-reported and genetic sex.

Due to sibling relationships in the WRAP cohort, genetic ancestry was assessed using Principal Components Analysis in Related Samples (PC-AiR), a method that makes robust inferences about population structure in the presence of relatedness (Conomos, Miller, & Thornton, 2015). This approach included several iterative steps and was based on 63,503 linkage disequilibrium (LD) pruned (r^2^<0.10) and common (MAF>0.05) variants, using the 1000 Genomes data as reference populations (Genomes Project et al., 2015). First, kinship coefficients (KCs) were calculated between all pairs of individuals using genomic data with the Kinship-based Inference for Gwas (KING)-robust method (Manichaikul et al., 2010). PC-AiR was used to perform principal components analysis (PCA) on the reference populations along with a subset of unrelated individuals identified by the KCs. Resulting principal components (PCs) were used to project PC values onto the remaining related individuals. All PCs were then used to recalculate the KCs taking ancestry into account using the PC-Relate method, which estimates KCs robust to population structure (Conomos, Reiner, Weir, & Thornton, 2016). PCA was performed again using the updated KCs, and KCs were also estimated again using updated PCs. The resulting PCs identified 1,198 WRAP participants whose genetic ancestry was primarily of European descent. This procedure was repeated within this subset of participants (excluding 1000 Genomes individuals) to obtain PC estimates used to adjust for population stratification in subsequent genomic analyses. Among European descendants, 160 variants were not in Hardy-Weinberg equilibrium (HWE) and 327,064 were monomorphic and thus, removed.

A total of 1,294,660 bi-allelic autosomal variants among 1,198 European descendants remained for imputation, which was performed with the Michigan Imputation Server v1.0.3 (Das et al., 2016), using the Haplotype Reference Consortium (HRC) v. r1.1 2016 (McCarthy et al., 2016) as the reference panel and Eagle2 v2.3 (Loh et al., 2016) for phasing. Prior to imputation, the HRC Imputation Checking Tool (Rayner, Robertson, Mahajan, & McCarthy, 2016) was used to identify variants that did not match those in HRC, were palindromic, differed in MAF>0.20, or that had non-matching alleles when compared to the same variant in HRC, leaving 898,220 for imputation. A total of 39,131,578 variants were imputed. Variants with a quality score R^2^<0.80, MAF<0.001, or that were out of HWE were excluded, leaving 10,400,394 imputed variants. These were combined with the genotyped variants, leading to 10,499,994 imputed and genotyped variants for analyses. Data cleaning and file preparation were completed using PLINK v1.9 (Chang et al., 2015) and VCFtools v0.1.14 (Danecek et al., 2011). Coordinates are based on GRCh37 assembly hg19.

## Author Contributions

B.F.D. conceptualization of the study, performed analyses, and wrote the manuscript; R.L.K., K.J.H, and S.C.J. provided critical manuscript revisions; and C.D.E. provided guidance for all aspects of the study.

## Acknowledgements

The authors thank the University of Wisconsin Madison Biotechnology Center Gene Expression Center for providing Illumina Infinium genotyping services. We especially thank the WRAP participants. The authors declare no conflicts of interest. BFD was supported by an NLM training grant to the Computation and Informatics in Biology and Medicine Training Program [NLM 5T15LM007359]. This research was also supported by the NIH [R01AG054047, R01AG27161, UL1TR000427, and P2C HD047873], Helen Bader Foundation, Northwestern Mutual Foundation, Extendicare Foundation, and State of Wisconsin.

